# Legacy Effects of Early β-Adrenergic Stimulation Program Adipose Plasticity and Confer Metabolic Resilience in Obesity

**DOI:** 10.64898/2026.06.23.734002

**Authors:** Pablo E. Morales, Wenxin Tong, Lavanya Vishvanath, Danielle C. Leander, Taylor E. Wade, David S. Hallaron, Kimberley El, Robert A. Hollander, Ashley Truong, Donald Wothe, George Elmquist, Marjori Russo, Heather K. Hamilos, Grace E. Dewyer, Clair Crewe, William L. Holland, Timothy R. Koves, Deborah M. Muoio, David A. D’Alessio, Jessica Cannavino, Jonathan E. Campbell, Mengle Shao, Rana K. Gupta

## Abstract

Pathologic white adipose tissue (WAT) remodeling, characterized by fibrosis, inflammation, and adipocyte dysfunction, is a hallmark and driver of metabolic disease in obesity^1^. Here, we show that legacy effects of early physiological or pharmacological interventions driving adaptive adipose remodeling can mitigate maladaptive WAT remodeling and metabolic dysfunction when developing obesity later in life. Cold exposure or β3-adrenergic receptor (β3AR) agonism (CL316,243) induced thermogenic remodeling of WAT in male mice. After a prolonged recovery at room temperature, trained epididymal WAT reverted to an energy-storing state but retained a population of adipocytes resembling “metabolically flexible” visceral adipocytes found in human metabolically healthy obesity. The legacy of the antecedent treatment conferred lasting protection against glucose intolerance when later developing high fat diet (HFD)-induced obesity, with insulin sensitivity persisting for at least 20 weeks of overnutrition. This metabolic resilience was accompanied by healthy epididymal WAT expansion with reduced fibrosis and inflammation. Our findings demonstrate that short-term interventions, without genetic manipulation, can “train” adipose tissue, enhancing its long-term plasticity and conferring durable protection against future obesity-associated insulin resistance.

## Main

Metabolic health varies widely among individuals with obesity, affecting risk of diabetes and response to emergent weight-loss therapies^2,3^. Comparisons of metabolically healthy obesity (MHO) to metabolically unhealthy obesity (MUO) highlight white adipose tissue (WAT) dysfunction, rather than mass alone, as a driver of metabolic disease^2,4,5^. WAT function in obesity is dependent on how it remodels as it expands. Healthy adaptive adipose remodeling includes angiogenesis and *de novo* adipocyte differentiation (“adipogenesis”), which sustain tissue growth and energy storage while preserving functional adipocytes^4,6^. Loss of this plasticity leads to maladaptive remodeling of expanding adipose tissue^4,7^. This pathologic remodeling is characterized by fibrosis, inflammation, and adipocyte dysfunction, and predicts insulin resistance and poor weight-loss response to bariatric surgery^8,9^. Although progress has been made in understanding the importance of WAT remodeling, the physiological or environmental determinants of these divergent trajectories remain poorly defined.

Our prior work showed that enhancing adipogenesis early in life has long-lasting effects on visceral epididymal WAT (eWAT) plasticity^10^. Transient overexpression of PPARγ, the “master regulator” of adipogenesis, in murine PDGFRβ+ progenitors only during the first week of life durably reprogrammed adipocyte progenitor cells well into adulthood. Although this intervention did not prevent diet-induced obesity, it protected against maladaptive eWAT remodeling and glucose intolerance. The findings from this genetically engineered model highlight how a limited intervention that drives adipose remodeling can influence the ability of the tissue to adapt to subsequent challenges later in life. Here, we asked whether physiological or pharmacological interventions, without genetic manipulation, can similarly train adipose plasticity and determine the metabolic outcome in future challenges to overnutrition.

The cold-induced transformation of WAT into an energy-burning brown fat-like phenotype (colloquially, “browning”) is exemplary of adaptive tissue remodeling. Cold exposure converts white adipose tissue into a more metabolically active, brown adipose tissue (BAT)-like state^11^. Local catecholamine release from sympathetic nerves stimulates adrenergic receptors on adipocytes, activating metabolic programs that drive lipid turnover, fatty acid oxidation, and heat production via UCP1-dependent and UCP1-independent ATP-consuming cycles^11,12^. The resulting “beige” adipocytes are activated alongside extensive tissue remodeling that includes angiogenesis, immune cell changes, and new beige adipocytes emerging from resident progenitors. Pharmacological agonism of the β3 adrenergic receptor (β3AR) (e.g., CL316,243), whose expression in mice is restricted to mature adipocytes, mimics many of the effects of cold exposure on adipose tissue^13,14^. This thermogenic state reverses when cold or β3AR stimulation ends, with “re-whitened” depots appearing morphologically indistinguishable from WAT that never experienced these interventions^15–17^. However, epigenomic and cellular changes persist in inguinal and brown adipocytes, creating a memory that primes future cold responses^16,18^.

We used cold exposure and β3AR agonism to test whether brief, intense remodeling can produce lasting benefits for adipose plasticity. In our experimental design (Fig. 1a), 6-week-old male C57BL/6N mice were cold (6 °C) exposed (CE) or treated with CL316,243 (CL). This age was chosen since 1) major WAT depots, including eWAT, are fully developed by this age, and 2) the thermogenic remodeling induced by these stimuli is most robust in young-adult mice^19^. We exposed animals to cold or treated daily with CL316,243 for 2 weeks, with the intention of maximizing remodeling even in the typically browning-resistant eWAT. Mice were then left undisturbed at room temperature for 4 weeks to allow adipose “re-whitening” before subsequent metabolic challenges.

**Figure 1:**
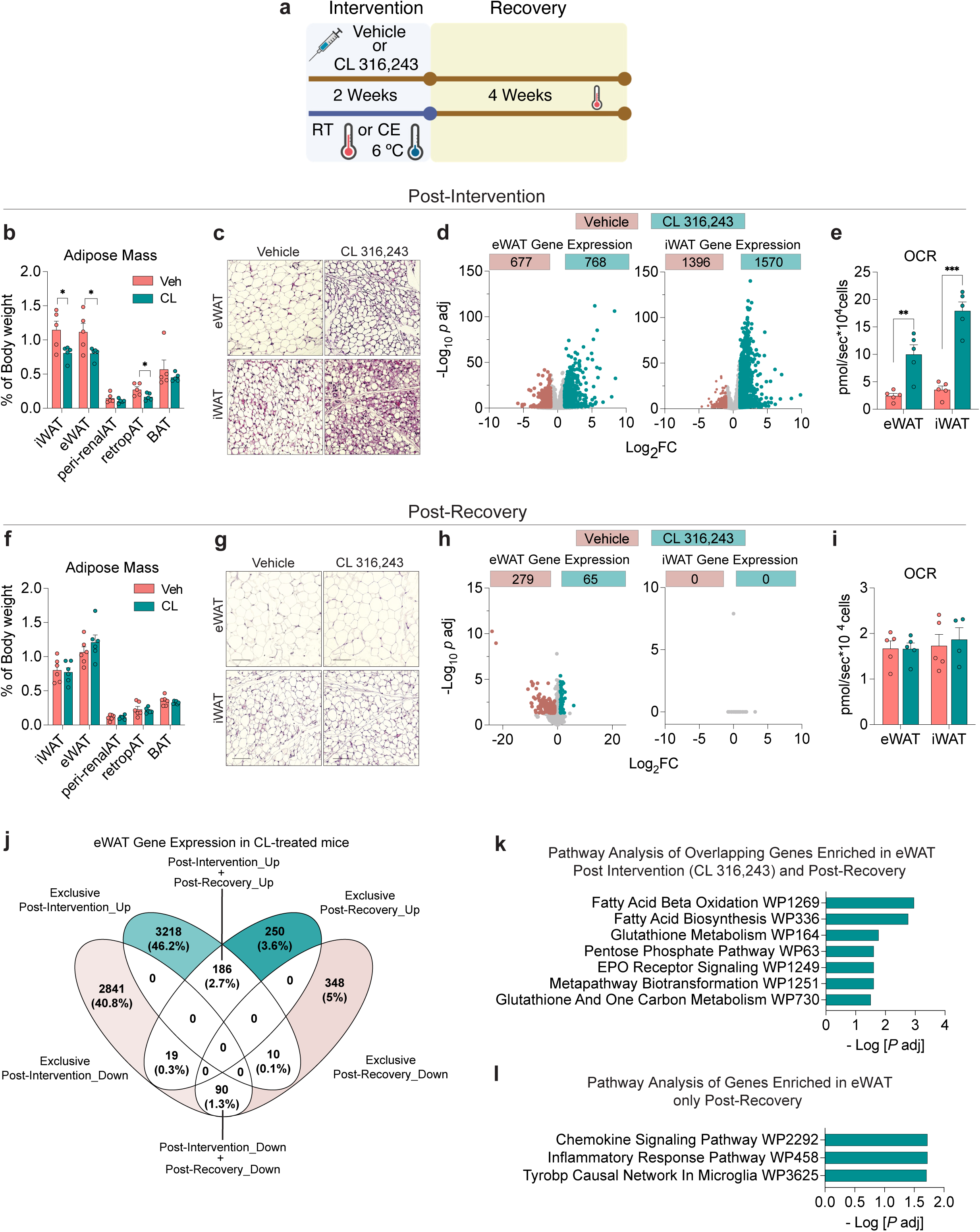
β3AR agonism and withdrawal remodel adipose tissue morphology and transcriptional programs. **a)** Experimental Design: 6-weeks-old male C57BL/6N mice were either 1) single-housed at 6°C (cold exposure, CE) or at room temperature (RT), or 2) treated daily with the β3AR agonist CL 316,243 (CL) (1 mg/kg body weight) or saline (vehicle; Veh) at RT for 2 weeks. Following the 2-week intervention period, animals were returned to RT housing for 4 weeks (Recovery). Image created in BioRender. **b)** Adipose depot mass following 2 weeks of vehicle or CL 316,243 treatment. n= 5-6. * denotes p <0.05 by unpaired t-test. **c)** Representative images of H&E-stained epididymal WAT (eWAT) and inguinal WAT (iWAT) from animals harvested following 2 weeks of vehicle or CL 316,243 treatment. Scale bar = 100 μm. **d)** Volcano plots depicting number of differentially expressed genes (p_adj_ ≤0.05; absolute log_2_FC ≥1) in whole eWAT or iWAT following 2 weeks of CL 316,243. Numbers of genes enriched in vehicle (negative fold change) or CL 316,243 (positive fold change) groups are shown. **e)** Routine oxygen consumption rate measured from primary fractionated adipocytes collected following 2 weeks of CL 316,243. Bars represent mean + SEM. Each data point represents a unique animal. n= 4-5. * denotes p <0.05 by unpaired t-test. **f-i)** Same as panels b-e, except 4 weeks after cessation of CL 316,243 treatment. **j)** Venn diagram depicting common and distinct patterns of eWAT gene expression differences following CL 316,243 treatment (Post-Intervention) and following recovery (Post-Recovery). **k)** Pathway Analysis of 186 genes induced in eWAT by CL 316,243 (p_adj_. p<0.05) and remain elevated following recovery. Overrepresented WikiPathways (p_adj_. p<0.05) are shown. **l)** Pathway Analysis of 250 genes induced in eWAT only during the recovery from CL 316,243 (p_adj_. p<0.05). Overrepresented WikiPathways (p_adj_. p<0.05) are shown. For all panels, bars represent mean + SEM. Each data point represents a unique animal. Exact measurements and statistical data for panels b, e, f, and i are provided in Source Data Figure 1. Full datasets for panels d,h, j-l, are in Supplementary Tables 1,2,4,5, 9-12.

After intervention, eWAT mass was slightly reduced in cold-exposed mice (Extended Data Fig. 1a), and inguinal WAT (iWAT), and retroperitoneal WAT were modestly smaller in CL-treated mice (Fig. 1b). Both stimuli produced multilocular adipocytes with the expected depot-specific differences (Fig. 1c; Extended Data Fig. 1b). Two weeks of CL316,243 altered hundreds of genes (padj < 0.05, >2-fold) across WAT and BAT and increased adipocyte respiration (Fig. 1d,e; Extended Data Fig. 1c; Supplementary Tables 1-3). After four weeks of recovery, adipose mass, morphology, global gene expression, and respiration returned to levels indistinguishable from controls (Fig. 1f-i; Extended Data Fig. 1a-c; Supplementary Tables 4-6).

In comparison to iWAT and interscapular BAT, more differentially expressed genes were found in eWAT following recovery from CL treatment (Fig. 1h, Extended Data Fig. 1c). Pathways enriched by CL treatment involved mitochondrial metabolism and the TCA cycle, consistent with beige adipocyte formation (Extended Data Fig. 1d; Supplementary Tables 7). After recovery, pathways related to inflammatory responses were overrepresented (Extended Data Fig. 1e; Supplementary Tables 8). Examining all genes significantly altered by CL intervention and/or recovery (padj < 0.05, any fold-change), we identified 186 transcripts persistently enriched after CL (Fig. 1j, Supplementary Table 9) and 250 newly enriched only after recovery (Fig. 1j, Supplementary Table 11). Persistent genes include adipocyte-enriched genes linked to fatty acid oxidation and synthesis (e.g., *Hadha*, *Acss2*, *Acaca*) (Fig. 1k; Supplementary Table 10), whereas recovery-specific genes were enriched for inflammatory signaling (Fig. 1l; Supplementary Table 12).

We reasoned that the subtle differences in gene expression revealed through bulk-RNA sequencing of whole tissue may reflect activity occurring within specific cell types or even cell subpopulations. Thus, we profiled CL-trained eWAT post-recovery at higher cellular resolution through single-cell (sc) and single-nuclei (sn) RNA sequencing (RNA-seq). scRNA-seq of CD45⁺ immune cells revealed the major immune populations in both groups (Extended Data Fig. 2a,b; Supplementary Table 13). Unsupervised clustering identified expected macrophage subsets, including perivascular (MC1, MC2; *Lyve1*⁺), non-perivascular (MC4; *Ear2*⁺), and cycling macrophages (MC3, MC6; *Mki67*⁺) (Extended Data Fig. 3a,b; Supplementary Table 14). Flow cytometry analyses indicated that the frequencies of total and CD163⁺ perivascular macrophages in eWAT were comparable between control and CL-trained mice (Extended Data Fig. 3c; Supplementary Figure 1). However, transcriptional differences emerged within specific macrophage subpopulations (Supplementary Tables 15–28). Genes enriched in several macrophage clusters of CL-trained eWAT mapped to phagocytic and efferocytosis pathways (e.g., *Trem2*, *Lipa*, *Axl*, *Tyrobp*) and included markers of lipid associated macrophages (LAMs) (Extended Data Fig. 3d). LAMs are phagocytic macrophages that localize to crown-like immune structures in WAT, where they remove dying dysfunctional adipocytes and mediate lipid recycling^20,21^. The frequency of eWAT LAMs is low in lean mice but then increases with diet-induced obesity to become the predominant macrophage population^22^. The adoption of a phagocytic phenotype after the cessation of CL316,243 may simply reflect a similar purpose: the need to remove remnant cells/debris and turn over lipid as tissues revert to an energy-storing WAT phenotype.

Our observation that CL-induced genes linked to fatty acid metabolism remain enriched during the recovery period motivated a focused higher resolution analysis of mature adipocytes. We conducted snRNA-seq, identifying adipocytes and the expected stromal-vascular cell types in both vehicle and CL-trained eWAT (Extended Data Fig. 2c,d; Supplementary Table 29). Unsupervised subclustering of adipocytes suggested potential differences in adipocyte gene expression. Adipocytes subclustered into three molecularly distinct cellular populations (Ad1, Ad2, Ad3) that were present in both control and CL-trained eWAT (Fig. 2a; Supplementary Table 30). Ad1 is marked by *Nnat* expression (*Nnat*^High^), enriched for genes related to ECM remodeling and growth factor signaling (Fig. 2b,c; Supplementary Table 31). The defining markers of Ad2 included genes related to core adipocyte functions such as triglyceride synthesis and fatty acid oxidation (Fig. 2b,d; Supplementary Table 32). Ad3 is a small cluster enriched for genes encoding ribosomal proteins, translation factors, and mRNA processing genes (Fig. 2b,e; Supplementary Table 33). Notably, Ad2 appeared enriched in CL-trained eWAT following recovery (Fig. 2a). Using *EnrichR*^23^, we asked whether the defining Ad2 gene signature resembled any annotated experimentally defined signatures of transcription factor perturbations. We found that gene signatures consistent with elevated EBF1, PPARγ, and PRDM16 activity are overrepresented in Ad2 (Fig. 2f; Supplementary Table 34). While mRNA levels of *Ebf1*, *Ebf2*, and *Pparg* are not enriched in Ad2 (Fig. 2g), shared downstream target genes that are regulated by these transcription factors are enriched, including key regulators of fatty acid uptake (*Slc27a1*, *Scarb1*), triglyceride synthesis (*Lpin1*), and lipolysis (*Abhd5*, *Pnpla2*) (Fig. 2h). Moreover, genes encoding known secreted proteins regulating adipogenesis (*Fgf10*) and angiogenesis (*Vegfa*) are enriched in Ad2. This subpopulation of adipocytes does not appear to be active beige adipocytes since classical beige markers, such as *Ucp1* and *Dio2*, were not detected. Thus, β3AR stimulation induces reversible remodeling that leaves persistent transcriptional imprints in adipocytes and distinct immune signatures during re-whitening.

**Figure 2:**
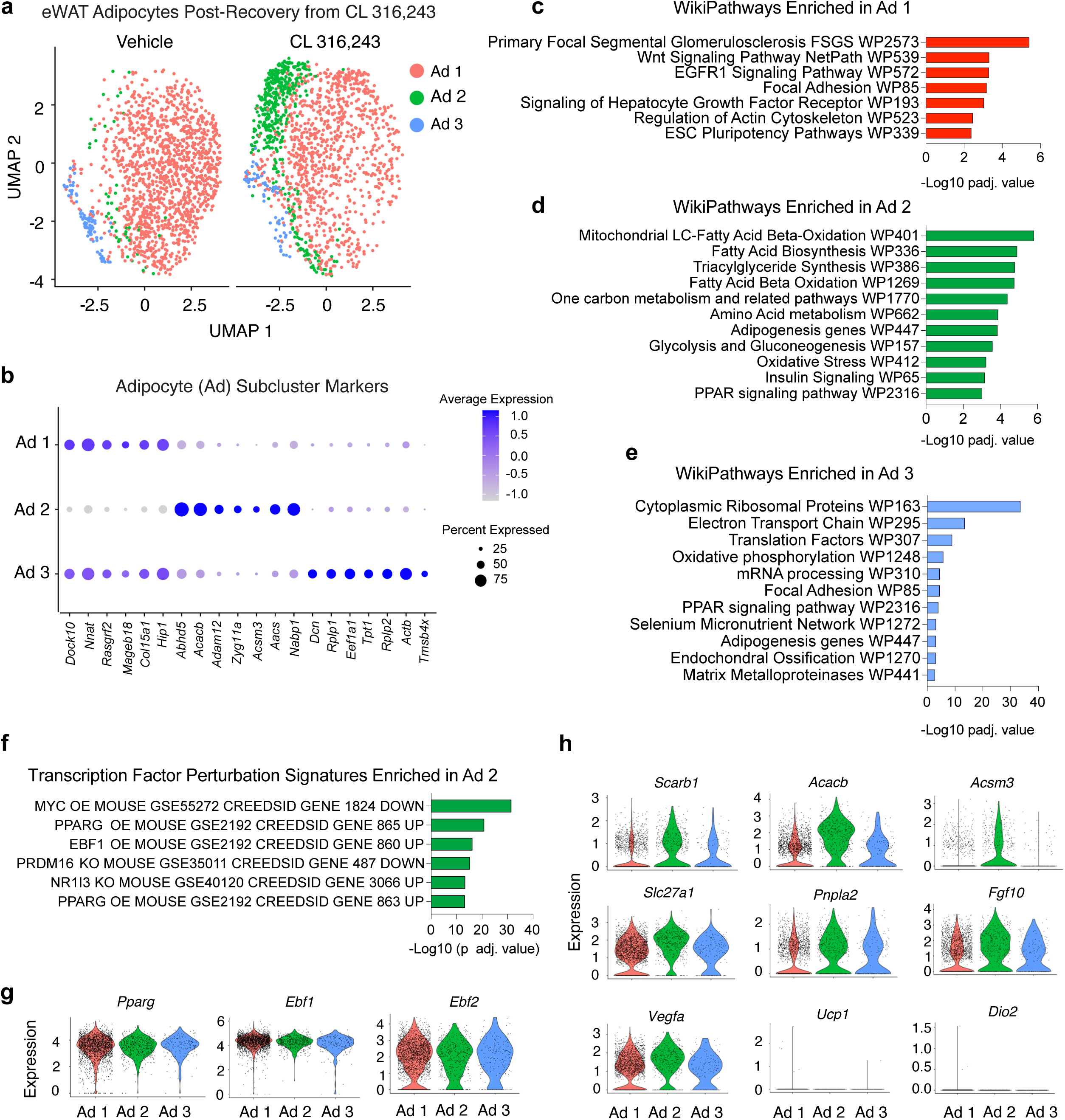
β3AR agonism induces durable transcriptional reprogramming in epididymal adipocytes. **a)** UMAP of single epididymal adipocyte nuclei transcriptomes obtained following the recovery from CL 316,243, revealing three molecularly distinct adipocyte (Ad) subclusters (Ad1-3). **b)** Heatmap depicting expression of most enriched transcripts defining Ad1-3. **c)** Overrepresented WikiPathways amongst genes enriched in Ad 1 **d)** same as panel c) for Ad 2 **e)** same as panel c) for Ad 3 **f)** Experimentally defined gene signatures of indicted transcription factor perturbations overrepresented in Ad 2. **g,h)** Violin plots of indicated gene expression in adipocyte subclusters. *EnrichR* was used for pathway analysis, with defining transcripts of each cluster (log_2_FC >0.25 and p_adj_. p<0.05) used as input. Full datasets for each panel are in Supplementary Tables 30-34.

We examined how cold or CL-trained mice under our paradigm adapt to overnutrition. We challenged male mice to a high-fat diet (HFD) (60% kcal fat) for at least ten weeks, beginning immediately after the four-week recovery period (Fig. 3a). Cold- or CL-training did not impact diet induced weight gain (Fig. 3b,c), but notable effects on glucose homeostasis were observed. After ten weeks on HFD, cold-trained animals displayed slightly better glucose tolerance than controls; however, the results were variable amongst independent cohorts and often did not reach statistical significance (representative experiment shown in Fig. 3d). On the other hand, CL-trained mice were highly and reproducibly more glucose tolerant than control mice (Fig. 3e). In both trained groups, plasma insulin excursions after glucose challenge were lower, and adiponectin levels higher (Fig. 3f–i), indicating relatively preserved insulin sensitivity. Hyperinsulinemic-euglycemic clamps after 20 weeks of HFD confirmed sustained insulin sensitivity in CL-trained mice, even 24 weeks after treatment cessation (Fig. 3j).

**Figure 3:**
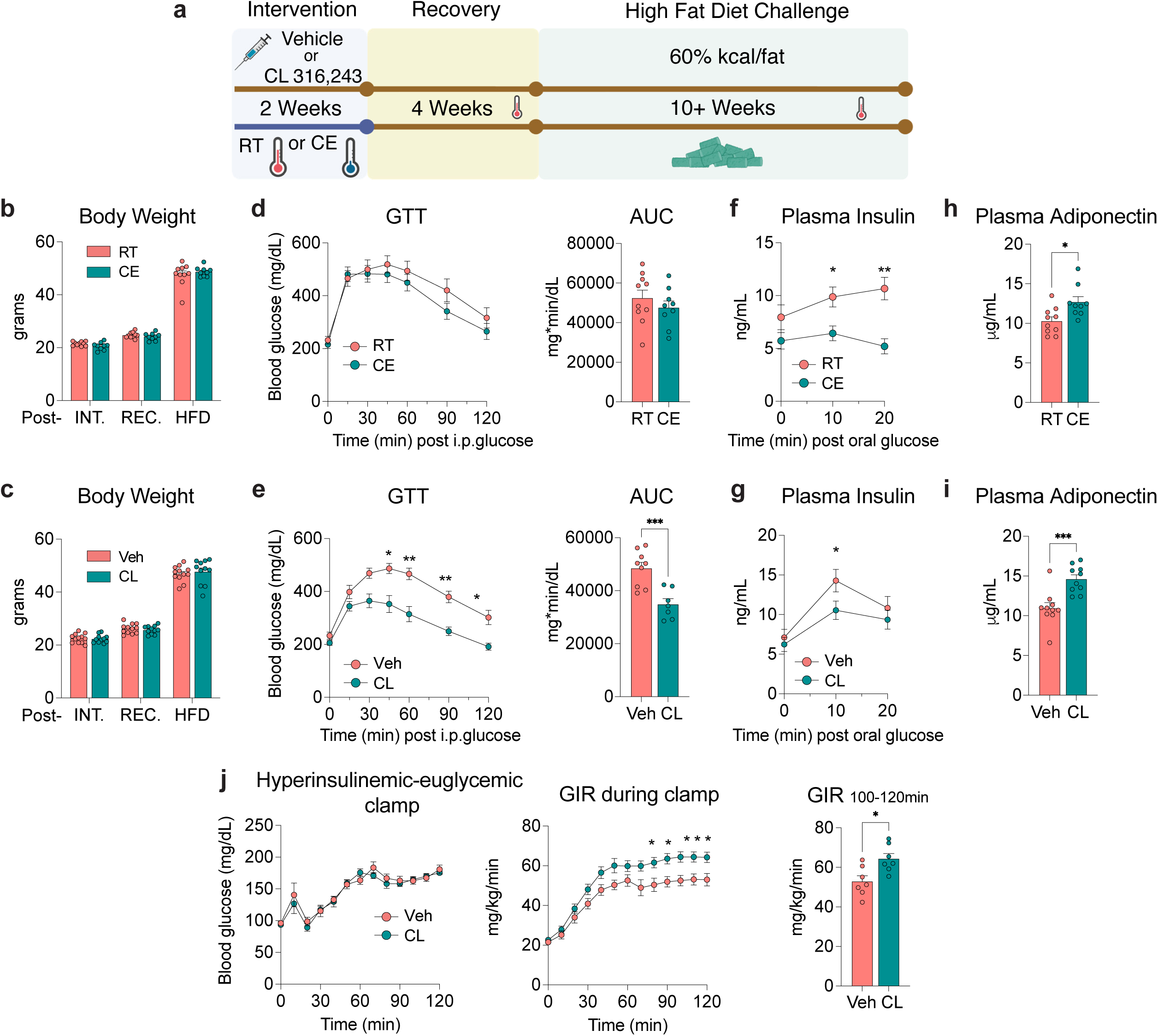
Early cold or β3AR training ameliorates insulin resistance during future obesity. **a)** Experimental Design: 4 weeks after cold exposure or CL316,243 treatment, animals were switched to a high fat diet (60% kcal from fat) and maintained at RT for ≥ 10 weeks (“HFD Challenge”). **b)** Average body weights of animals following the cold intervention (Post-INT), the recovery period (Post-REC), and after 12 weeks of HFD feeding (Post-HFD). n= 9-10. **c)** Same as panel B, for animals treated with vehicle or CL 316,243. n= 11-12. **d)** (left) Intraperitoneal glucose tolerance tests (GTT) of control (RT) or cold-trained mice (CE) after 11 weeks of HFD feeding. (right) Area under the GTT curve (AUC) measurements. n=9-10 per group. **e)** Same as panel D, for animals treated with vehicle or CL 316,243. n=7-9 per group. For GTT, * denotes p<0.05, ** p<0.01 by two-way ANOVA followed by Šídák’s multiple comparisons test. *** denotes p <0.001 by unpaired t-test test for AUC measurement. **f)** Plasma insulin levels in control and cold-trained mice after HFD feeding. Insulin measurements were made at indicated time points following single oral glucose bolus. n= 11 * denotes p <0.05, ** denotes p<0.01 by two-way ANOVA followed by Šídák’s multiple comparisons test. **g)** Same as panel F, for animals treated with vehicle or CL 316,243. n= 9-12. **h)** Plasma Adiponectin levels in control and cold-trained mice following 11 weeks of HFD. n= 9-10 per group. * denotes p <0.05 by unpaired t-test. **I)** Same as panel H, for animals treated with vehicle or CL 316,243. n= 10. *** denotes p <0.001 by unpaired t-test. **j)** Hyperinsulinemic-euglycemic clamps after 18 weeks of HFD feeding (n=7 mice per group): (left) Blood glucose levels during insulin infusions, (middle) Glucose infusion rate during insulin clamp, (right) Average glucose infusion rate over minutes 90-120. For GIR over time, * p <0.05 by two-way ANOVA followed by Šídák’s multiple comparisons test. For GIR_90-120_, * p <0.05 by unpaired t-test. Insulin clamps were performed in two different cohorts. Representative experiment is shown. For all panels, bars represent mean + SEM. Each data point represents a unique animal. Exact measurements and statistical data are provided in Source Data Figure 3.

We also assessed how prior cold- or CL-training impacted adipose tissue remodeling. After 12 weeks of HFD feeding, trained mice had visibly larger visceral eWAT depots and modestly smaller livers than controls animals, with no significant differences in the mass of other adipose tissues examined (Fig. 4a). Despite the larger size, eWAT from both cold- and CL-trained mice exhibit far fewer MAC2+ crown-like immune structures and less collagen deposition than in control eWAT, indicating protection from local inflammation and fibrosis (Fig. 4b,c). This was corroborated by gene expression analyses revealing lower transcript levels of collagens and immune cell markers (Extended Data Fig. 4a). Mean epididymal adipocyte size was not impacted by prior cold training (Fig. 4d). Surprisingly, mean epididymal adipocyte size was slightly larger in CL-trained animals (Fig. 4d). This was unexpected as countless cross-sectional analyses of human WAT implicate adipocyte hypertrophy as a feature of pathologic WAT remodeling that correlates with insulin resistance. However, this result is not unprecedented and echoes the adipose phenotype of previously described *Col6* deficient mice with hypertrophic WAT expansion associated with metabolic protection^24^. It is postulated that weakening of the extracellular matrix allows for stress-free adipocyte expansion^24^. Moreover, longitudinal overfeeding studies do not support a clear-cut relationship between adipocyte size and metabolic health^25^. Nevertheless, we performed parallel lineage tracing studies to evaluate whether eWAT expansion in cold- and/or CL-trained mice also includes cellular hyperplasia via adipogenesis. We utilized the previously described doxycycline (DOX)-inducible *Pdgfrb*-lineage tracing system, which has been used to track for the formation of new adipocytes formed during HFD feeding in adult mice (Extended Data Fig. 4b)^7,26,27^. We induced indelible labeling of progenitor cells with membrane GFP (mGFP) expression during the last week of the 4-week recovery from either cold or CL 316,243 treatment. We then placed mice on HFD for 10 weeks without doxycycline (Extended Data Fig. 4c). Indeed, mGFP+ adipocytes, indicative of newly formed fat cells, were more readily observed in cross sections of eWAT, but not iWAT, in trained animals (Extended Data Fig. 4d). These data indicate that the degree of *de novo* adipogenesis that occurs in association with overnutrition can be enhanced by durable effects of prior interventions, and in a depot-specific manner.

**Figure 4:**
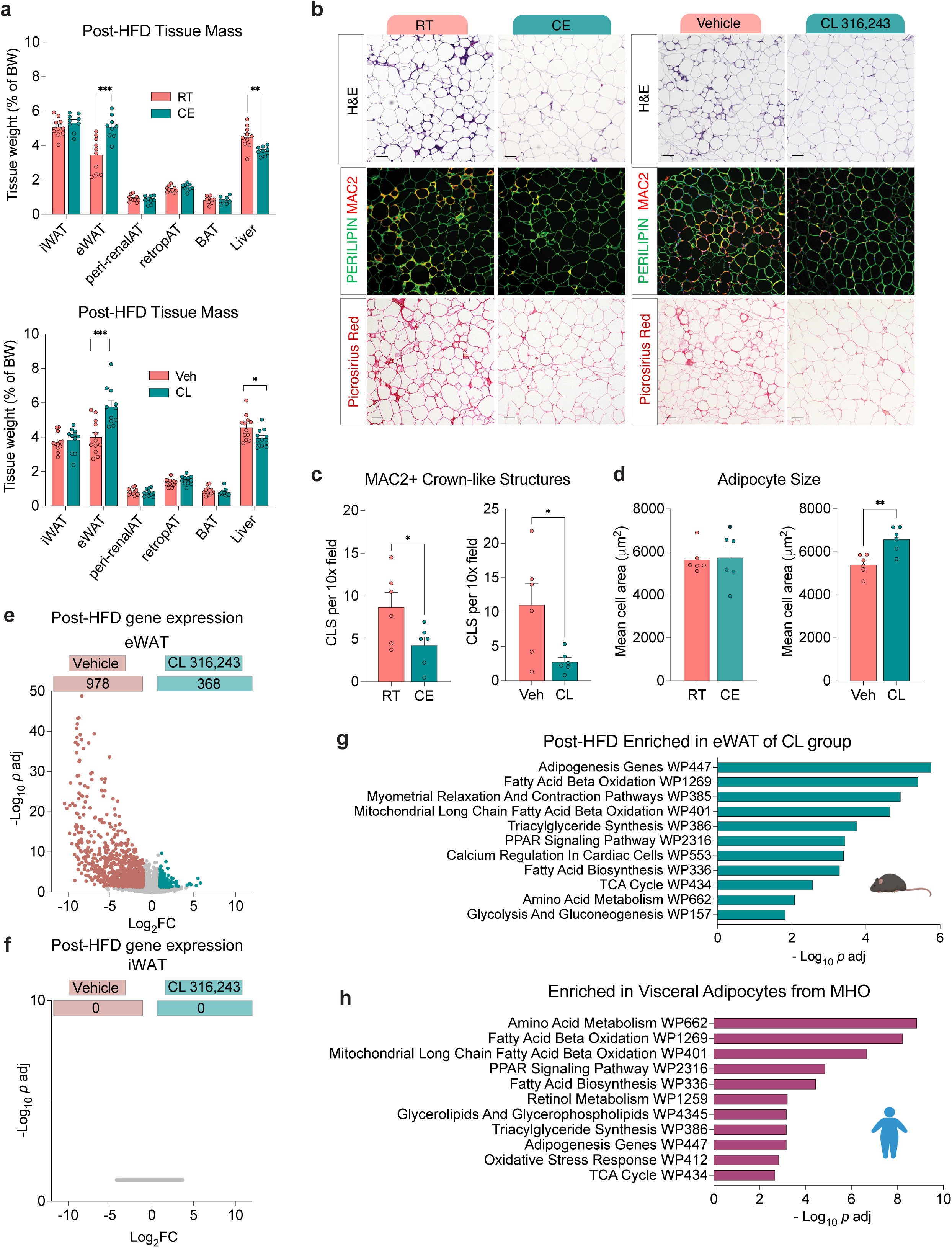
Early cold or β3AR training promotes adaptive WAT expansion during future obesity. **a)** Mass of indicated tissues in cold-trained (top) or CL-trained mice (bottom) after 12 weeks of HFD feeding. Top, n= 9-10; bottom, n = 11-12. * denotes p<0.05, ** p<0.01, *** p<0.001 by unpaired t-test. **b)** Histological analyses of eWAT from cold-trained (left) or CL-trained (right) animals after 12 weeks of HFD feeding. Top row: hematoxylin and eosin (H&E) staining; scale bar = 200 μm. Middle row: indirect immunofluorescence of PERILIPIN and MAC2 expression. Bottom row: picrosirius red staining of collagen deposition. scale bar = 200μm. **c)** Frequency of MAC2+ crown-like immune structures (CLS) in eWAT of cold-trained (left) or CL-trained (right) after 12 weeks of HFD feeding. n=6 mice per group. * p<0.05 by unpaired t-test. **d)** Mean adipocyte size in eWAT of cold-trained (left) or CL-trained (right) after 12 weeks of HFD feeding. n=6 mice per group. ** denotes p<0.01 by unpaired t-test. **e)** Volcano plot depicting differentially expressed genes (p_adj_ ≤0.05; absolute log_2_FC ≥1) in eWAT of CL-trained mice after 12 weeks of HFD feeding. n=4-6 mice for each group. Numbers of genes enriched in vehicle (negative fold change) or CL 316,243 (positive fold change) groups are shown. **f)** same as panel e) for iWAT **g)** Top 11 annotated WikiPathways enriched (p adj. <0.05) in eWAT of CL 316,243 treated mice (post-HFD). **h)** Top 11 annotated WikiPathways enriched (p adj. <0.05) in visceral adipocytes of individuals with MHO compared to BMI-matched individuals with MUO. EnrichR analysis was performed using list of differentially expressed genes (MHO vs. MUO) described in Reinisch et al. For panels a, c, d, and e, bars represent mean + SEM. Each data point represents a unique animal. Complete EnrichR analysis for panels g,h can be found in Supplementary Tables 35,36.

We performed bulk-RNA sequencing of WAT from vehicle and CL-trained mice after 12 weeks of HFD feeding to gain molecular insight into the adipose phenotype. We observed robust alterations in eWAT global gene expression (Fig. 4e; Supplementary Table 35). The same analysis parameters did not reveal any statistically significant differences in iWAT gene expression (Fig. 4f; Supplementary Table 36). Pathway analysis of genes statistically enriched in eWAT of CL-trained mice with obesity revealed an overrepresentation of genes driving lipid handling (triglyceride synthesis, fatty acid oxidation), adipogenesis, and PPARγ signaling (Fig. 4g; Supplementary Table 37), mirroring pathways defining the Ad2 adipocytes post-recovery (Fig. 2d). Moreover, the same gene signatures reflective of elevated EBF1, PPARγ, and PRDM16 activity in Ad2 were also overrepresented in eWAT of CL-trained mice after HFD feeding (Extended Data Fig. 5a; Supplementary Table 38). The enrichment of metabolic pathways in these CL-trained eWAT depots is also notable considering recent comprehensive analyses of WAT from MHO and MUO. Reinisch et al. reported that omental adipocytes from individuals with MHO are enriched in gene programs encoding effectors of triglyceride biosynthesis, lipid catabolism, and mitochondrial function^28^. The authors described these adipocytes as likely retaining “metabolic flexibility”, with adequate capacity for regulated lipogenesis, lipolysis, and adipokine production. Efthymiou et al. confirm and extend these findings, revealing that adipocyte subsets enriched gene signatures reflecting heightened adipocyte maturation and metabolic activity define WAT of MHO^29^. We conducted pathway analyses of the genes reported in Reinisch et al. that are significantly enriched in visceral adipocytes of MHO. Strikingly, the most overrepresented functional WikiPathways (Fig. 4h; Supplementary Table 39) and transcription factor perturbation signatures (Extended Data Fig. 5b; Supplementary Table 40) were the same as those most enriched in the eWAT of CL-trained mice exhibiting protection from insulin resistance in obesity.

Taken together, these data show that cold exposure or pharmacological agonism can durably program adipose plasticity, training WAT for an adaptive, instead of maladaptive, response to overnutrition faced later in life. These effects were strong enough to confer durable protection against insulin resistance for at least 20 weeks of HFD feeding, despite developing obesity. This phenomenon can be described as a “legacy effect”, defined in the context of medicine as “the continuous benefit of an intervention long after its cessation”^30^. From a teleological perspective, cold-induced “priming” events that ensure a metabolically healthy adaptation to future chronic overnutrition could serve hibernating animals as they transition from dormancy to the active feeding season. Our results build upon prior work showing that cold exposure can train future cold responses, mediated by memory in iWAT, BAT, or even germline epigenomic marks^16,18,31,32^. In humans, the precise relationship between environmental temperature and long-term metabolic health is still unclear but evidence supports an impact of cold exposure or pharmacological β3AR agonism on glucose homeostasis that goes beyond energy expenditure^33,34^. Of note, Speakman and colleagues found that the prevalence of type 2 diabetes within the United States, independent of obesity, poverty, and race, is lower in regions of lower ambient temperature^35^. Nevertheless, the severity of maladaptive WAT remodeling in obesity may also be mitigated by other physiological or pharmacological interventions that promote adaptive remodeling. Similar adipose “training” effects may arise from other stimuli invoking adrenergic signaling or cAMP-driven responses. Exercise, for example, warrant investigations in light of the known legacy effects of exercise on cardiometabolic parameters^36^.

There are some limitations to our study that require consideration going forward. Adipose remodeling is sex and age dependent^19,37,38^. Here, interventions were applied to 6-week-old male mice. Understanding whether the observed legacy effects are influenced by sex and age can yield insight into the optimal conditions for adipose training. The current studies do not firmly establish the mechanisms underlying the legacy effects of these interventions. However, our transcriptional profiling of eWAT following the recovery from CL 316,243 treatment raises some new hypotheses. The presence of phagocytic macrophages prior to the onset of HFD feeding may endow WAT with the ability to rapidly clear dying or stressed cells during tissue expansion. Our adipocyte analysis raises the possibility that cold or pharmacological β3AR agonism induces long-term retention of metabolically resilient, or “flexible”, cells resembling those seen in MHO. It is notable that EBF, PRDM16, and PPARγ target genes are enriched in the unique Ad2 adipocytes. It is plausible that these well-known transcriptional drivers of beige adipocyte activity also establish lasting transcriptional effects preserving adipocyte fitness after the stimulus is gone. Although speculative, this hypothesis aligns with the observations that EBFs and PRDM16 regulate adipose tissue metabolism through mechanisms beyond adaptive thermogenesis. Loss of *Zfp423*, a potent EBF repressor, in intra-abdominal WAT depots protects against pathologic WAT remodeling and glucose intolerance when developing obesity^39^. Similarly, adipocyte-specific PRDM16 overexpression protects against glucose intolerance independent of its impact on body weight^40^. EBF1 regulates adipocyte lipid handling and inflammatory programs ^41,42^. Importantly, levels of adipose EBF1 expression in human obesity correlate with metabolic health ^43^. Of course, it is also possible that the memory of these interventions is encoded in other cell types affecting tissue remodeling, including vascular cells, adipose progenitors, T cells, and neurons.

Our work establishes an experimentally tractable model of metabolically healthy WAT expansion without genetic manipulation, enabling dissection of such cellular and molecular determinants of adaptive remodeling and tissue memory. Importantly, this model also offers an experimental tool to study the physiological consequences of healthy WAT expansion in obesity, including any effects that may extend beyond glucose homeostasis. Ultimately, this line of investigation can help determine mechanisms underlying key aspects of the metabolic dysfunction commonly seen in human obesity, define new biomarkers of adipose tissue health, and lay the groundwork for developing strategies to maintain or restore adipose plasticity, and thereby metabolic resilience, throughout adulthood.

## Methods

### Animals

Animal experiments were performed according to procedures approved by Institutional Animal Care and Use Committees at Duke University School of Medicine, UT Southwestern Medical Center, and Washington University of St. Louis School of Medicine. C57BL6/N mice (Charles River) were used throughout the study. *Pdgfrb*^rtTTA^; *TRE-Cre*; *Rosa26R*^mT/mG^ mice were maintained on a C57BL/6 background and were previously described^27^. Mice were maintained on a 12hr light/dark cycle (7AM/7PM) in a temperature-controlled barrier facility (room temperature, ∼22°C) and given free access to water and food. Mice were maintained on a standard rodent chow diet (PicoLab Rodent Diet 20, LabDiet for Duke University studies) from the time of weaning (4 weeks of age). High fat diet (60% kcal/fat) was obtained from Research Diets (D12492i). Doxycyline-containing chow diet (600 mg/kg) was obtained from Bio-Serve (#S4107). Mice undergoing fasting had free access to water. For cold exposure studies, animals were single housed in temperature-controlled chambers (Powers Scientific, Inc.) set to 6 °C (maintaining temperature between 6-7 °C). Controls for cold exposure studies were animals single housed at room temperature. CL 316,243 was delivered once daily by intraperitoneal injection (1 mg/kg) for 14 days. Control mice were injected with equivalent volumes of saline. Following cold or CL 316,243 treatments, animals were maintained at room temperature for four weeks before sacrifice or commencement of high fat diet feeding.

### Histology and indirect immunofluorescence

Dissected tissues were fixed in 4% paraformaldehyde-PBS pH 7.4 for approximately 24 hours. Tissues were then rinsed in PBS and stored in 50% ethanol before tissue processing, paraffin embedding, and sectioning to 4 μm thickness. Picrosirius Red staining was performed following manufacturer’s instructions (Abcam, ab246832). For indirect immunofluorescence assays, paraffin sections were dewaxed in xylene and gradually dehydrated in decreasing concentrations of ethanol. Slides were submerged in Buffer A and incubated in an antigen retriever (Retriever 2100, Aptum Biologics Ltd.) and allowed to cool to room temperature before proceeding. Sections were treated with FX signal enhancer for 30 minutes, rinsed, then blocked in 10% normal goat serum/PBS for 30 minutes-1 hour. Primary antibodies were incubated overnight at 4°C. Secondary antibodies were incubated for 1 hour at room temperature. Antibodies and concentrations used for immunofluorescence include: guinea pig anti-PERILIPIN 1:500 (Biosynth #20RPP004) and rabbit anti-MAC-2 1:500 (Cedarlane, Clone M3/38, #CL8942AP). AlexaFluor secondary antibodies were chosen against original primary antibody source animal. Slides were mounted with Prolong Gold or Prolong Diamond mounting media containing DAPI and sealed using clear nail polish. Bright-field and fluorescence images were acquired using a Keyence BZ-X710 microscope. Cell size was determined using the Adiposoft plug-in for Fiji.

### Respirometry of isolated adipocytes

Isolated adipocytes were isolated as previously described^44^. Oxygen consumption was measured with the OROBOROS Oxygraph-2k (O2K) system. Assays were conducted in DMEM/F12 supplemented with 10% FBS as supporting buffer at 37°C with continuous stirring. Chambers were equilibrated with atmospheric oxygen before adding isolated adipocytes. Respiration from around 150,000 adipocytes from eWAT or iWAT, and 50,000 adipocytes from BAT was recorded until flux stabilization and interpreted as routine respiration.

### Glucose tolerance tests

For intraperitoneal glucose tolerance tests, animals were fasted for 6 hours and then injected with 1g/kg body weight of D-glucose (G7528, Sigma-Aldrich). Blood glucose was determined using Bayer Contour glucometers from tail blood.

### Insulin and Adiponectin measurements

Insulin measurements were performed following oral glucose challenge. After 5-6 hour fast, mice were administered 1.5g/kg body weight of D-glucose (G7528, Sigma-Aldrich) via oral gavage. Blood was collected from the tail before (0 min) and 10 and 20 minutes post-glucose bolus. Adiponectin measurements were obtained from blood collected immediately after 5-6 hour fast. Plasma was obtained by centrifugation at 10.000 x g at 4°C for 20 minutes and supernatant was frozen at −80°C until further use. Samples were thawed on ice and assayed according to the respective manufacturer’s instructions. Insulin (#10-1247-10, Mercodia) and Adiponectin (EZMADP-60k, Millipore) were measured by ELISA.

### Hyperinsulinemic Euglycemic Clamps

Animals were transferred to the clamp room 5 hours prior to the start of the clamp for fasting and acclimation to the room. Baseline samples were collected for glycemia, and then animals were connected to a tether-swivel system (Instech Laboratories Inc., PA). Syringe pumps with Insulin (4mU/kg/min) and donor red blood cells (1.375 uL/min) as well as 50% Dextrose at a variable rate were started after baseline. Samples were collected at 10-minute intervals to measure glycemia and to adjust the glucose infusion rate (GIR) to target euglycemia of 160-180 mg/dL. Steady state was reached 90 minutes into the clamp period, and whole blood samples were collected from 90 to 120-minutes for further analysis.

### Tissue RNA extraction

After dissection, tissues were snap-frozen in liquid nitrogen and kept at −80°C until further use. Samples were pulverized using a cryogenic tissue crusher in dry ice. RNA was extracted using columns (RNeasy Mini Kit, with DNAse digestion, #74104, Qiagen) with some modifications. Briefly, 50-100mg of powdered tissue was homogenized with 1mL of TRIzol reagent in a Tissue Lyser II (Qiagen) for 3 min. After a short centrifugation for discarding the top fat layer, 0.2mL of chloroform was added to the mix and centrifuged for 15 min at 12.000 xg at 4°C. The aqueous phase containing the RNA was recovered and then treated for the extraction using the affinity columns following the manufacturer’s instructions.

### Bulk mRNA sequencing and Data Analysis

mRNA libraries were prepared using the NEBNext Poly(A) mRNA Magnetic Isolation Module (New England Biolabs, E7490) and the NEBNext Ultra II RNA Library Prep Kit for Illumina (New England Biolabs, E7770). Indices used were from NEBNext Multiplex Oligos for Illumina (Index Primers Set 1, 2, 3 and 4, New England Biolabs). Sequencing was performed in a Novaseq 6000 instrument by the Sequencing and Genomic Technologies Shared Resource from the Duke University School of Medicine. FASTQ files were aligned to the mouse reference genome GRCm39 by STAR^45^ (Version 2.7.11b) excluding multiple mapped reads. The output BAM files were further counted using FeatureCounts^46^ (Version 1.6.3) for the raw count matrix generation. Raw counts were first normalized before performing the differential expression analyses using DESeq2^47^ (Version 1.44.0).Genes showing altered expression with adjusted p-value < 0.05 and |log2 FC| > 1 were considered statistical significant.

### Single-cell and single-nuclei RNA libraries

For nuclei isolation, eWAT stored at −80°C was pulverized using a cryogenic tissue crusher in dry ice and then homogenized with a Douncer homogenizer as previously described^48^. Nuclei from four different mice per condition were independently isolated and then pooled together for library preparation. Libraries were generated following the manufacturer’s instructions (10X Genomics, Pleasanton, CA, United States) and sequenced on a Novaseq X Plus instrument at the GCB Sequencing and Genomic Technologies Shared Resource, Duke University.

### Single-cell and single-nuclei RNA sequencing data analysis

The raw FASTQ files were processed using the Cell Ranger^49^ (Version 9.0.0) pipeline via the 10X Genomics Cloud for sample demultiplexing, read alignment to reference mouse genome GRCm39, cell barcode processing and unique molecular identifier (UMI) counting. All data were then loaded on R (Version 4.4.3) and first de-contaminated using SoupX^50^ (Version 1.6.2) before being read into Seurat^51^ (Version 4.4.3) for further analysis. For quality control, cell/nuclei with unique feature counts < 250, UMI counts < 500 and mitochondrial gene counts > 10% were discarded from downstream analyses. After filtration, only genes expressed in at least 2 cells were kept.

For each dataset (whole eWAT snRNAseq and CD45+ scRNAseq), the pooled “vehicle” and “CL 316,243” samples were normalized via using the function ScaleData and then integrated using canonical correlation analysis method of Seurat. The first 50 principal components of the PCA were computed using RunPCA before clustering was performed using FindNeighbors. The robustness of the clustering was assessed using clustree displaying the relationship between the clusters with increasing resolution. The marker genes of each cluster were computed using the FindAllMarkers function of Seurat for the selected clustering using the criteria of log2FC > log2(0.25) and a minimum fraction of 0.25 in all clusters for both the Likelihood-ratio test (test.use = “bimod”) and Wilcoxon Rank Sum test (test.use = “wilcox”). Clusters were then annotated into broad cell types based on known markers from literature. For differential gene expression analysis between vehicle and CL 316,243, counts were first added before comparing between conditions using NOISeq^52^ (Version 2.48.0). WikiPathways enrichment of the differentially expressed genes was performed using the R package enrichR^23^ (Version 3.4). The cell-cell interaction analysis in the snRNA samples was analyzed using the R package CellChat^53^ (Version 1.6.51). Adipocytes and macrophages were selected for each analysis and re-clustered following the same pipeline as mentioned above to identify the different cellular subtypes using the first 15 PCs. The marker genes of each sub-cluster were computed using the FindAllMarkers function of Seurat for the selected clustering using the criteria of log2FC > 0.25 and a minimum fraction of 0.25 in all clusters for both the Likelihood-ratio test (test.use = “bimod”) and Wilcoxon Rank Sum test (test.use = “wilcox”). Subclusters were manually annotated based on the top marker genes. For differential gene expression analysis between vehicle and CL 316,243, counts were first added before comparing between conditions using NOISeq (Version 2.48.0). WikiPathways enrichment of the marker genes was performed for each adipocyte subpopulation using the R package enrichR (Version 3.4)

## Data and code availability

Original data are provided as Source Data files or Supplementary Tables corresponding to each main and extended figure. Raw sequencing files will be deposited to Gene Expression Omnibus (GEO) upon re-opening of the NIH/United States government. Seurat objects of scRNA- and snRNA-seq studies and the associated data analysis pipelines for data integration, clustering, and secondary analyses, can be obtained from GitLab (https://gitlab.oit.duke.edu/morales-et-al/wat_cold_legacy_gupta). The scRNA- and snRNA-seq analyses can be further explored at https://guptalab-dmpi.shinyapps.io/shinyAppForPaper/.

## Acknowledgements

The authors are grateful to current and former members of the Gupta laboratory and Duke Molecular Physiology Institute for useful discussions. We thank the Duke University School of Medicine for the use of the Sequencing and Genomic Technologies Shared Resource, which performed all library sequencing. The authors appreciate the valuable collaboration and service of Molecular Genomics Core at the Duke Molecular Physiology Institute, which provided access to critical instrumentation and provided support for transcriptomics analyses. This study and personnel were supported, in part, by the National Institutes of Health, awards DK143978 and DK104789 to R.K.G., DK143978, DK123075, DK125353, and DK046492 to J.E.C., DK142423 to D.A.D., DK132461 to K.E., DK136168 to A.T., DK108833 and DK112826 to W.L.H., EB035738, DK137791, and DK020579 (Washington University Diabetes Research Center) to C.C., DK124723 (North Carolina Diabetes Research Center) to J.C., the American Diabetes Association Postdoctoral Fellowship Award 1-25-PDF-121 to W.T., the Career Development Award 19CDA34670007 from the American Heart Association and the Harry S. Moss Heart Trust to M.S, and the Duke University Borden Scholar Award to P.E.M.

## Author Contributions

Conceptualization: P.E.M., M.S., and R.K.G.; Validation (independent replication of key results): M.R. and C.C.; Formal analysis: P.E.M., M.S., W.T., J.C., T.E.W., D.L., C.C., W.L.H., J.E.C., and R.K.G.; Investigation: P.E.M., M.S., W.T., J.C., L.V., D.L., D.S.H., K.E., R.A.H., A.T., D.W., G.E., M.R., H.K.H., and G.E.D.; Critical resources and method development: T.R.K. and D.M.M.; Data Curation: P.E.M., M.S., W.T., T.E.W.; Writing - Original Draft: P.E.M., W.T., and R.K.G.; Writing - Review & Editing: all authors; Visualization: P.E.M., W.T., and R.K.G., Supervision: R.K.G., M.S., J.C., J.E.C., D.M.M., D.A.D., W.L.H., and C.C.; Project administration: R.K.G.; Funding acquisition: R.K.G., M.S., J.C., J.E.C., W.L.H., and C.C.

---

**Extended Data Figure 1:**
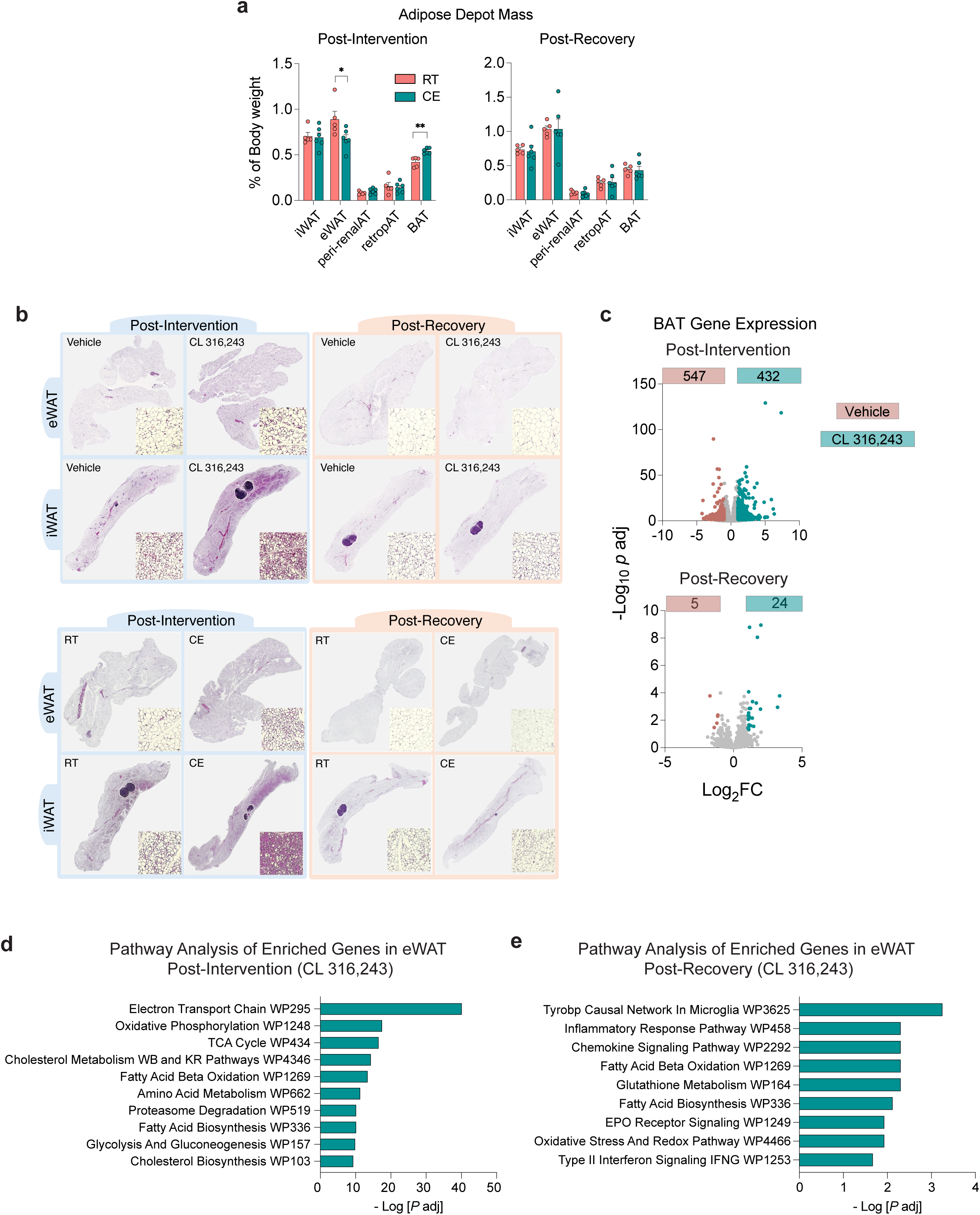
“Browning” and “Re-whitening” of Adipose Tissues. **a)** Adipose depot mass following 2 weeks of cold exposure (CE) (Post-Intervention, left) and after the 4-week recovery period (Post-Recovery, right). n= 5-6. * denotes p <0.05, ** denotes p <0.01 by unpaired t-test. Bars represent mean + SEM. Each data point represents a unique animal. Exact measurements and statistical data are provided in Source Data Extended Data Figure 1. **b)** Representative images of H&E-stained epididymal WAT (eWAT) and inguinal WAT (iWAT) following 2 weeks of the indicated intervention or after the 4-week recovery period at RT. **c)** Volcano plots depicting number of differentially expressed genes (p_adj_ ≤0.05; absolute log_2_FC ≥1) in interscapular BAT following 2 weeks of CL 316,243 (top) or after the recovery period (bottom). Numbers of genes enriched in vehicle (negative fold change) or CL 316,243 (positive fold change) groups are shown. **d)** EnrichR pathway analysis of genes enriched in eWAT (log_2_ FC >0.25; p_adj_ <0.05) after CL 316,243 treatment. **e)** Same as panel d, after CL 316,243 recovery. Full datasets for panels c-e are in Supplementary Tables 3, 6-8.

**Extended Data Figure 2:**
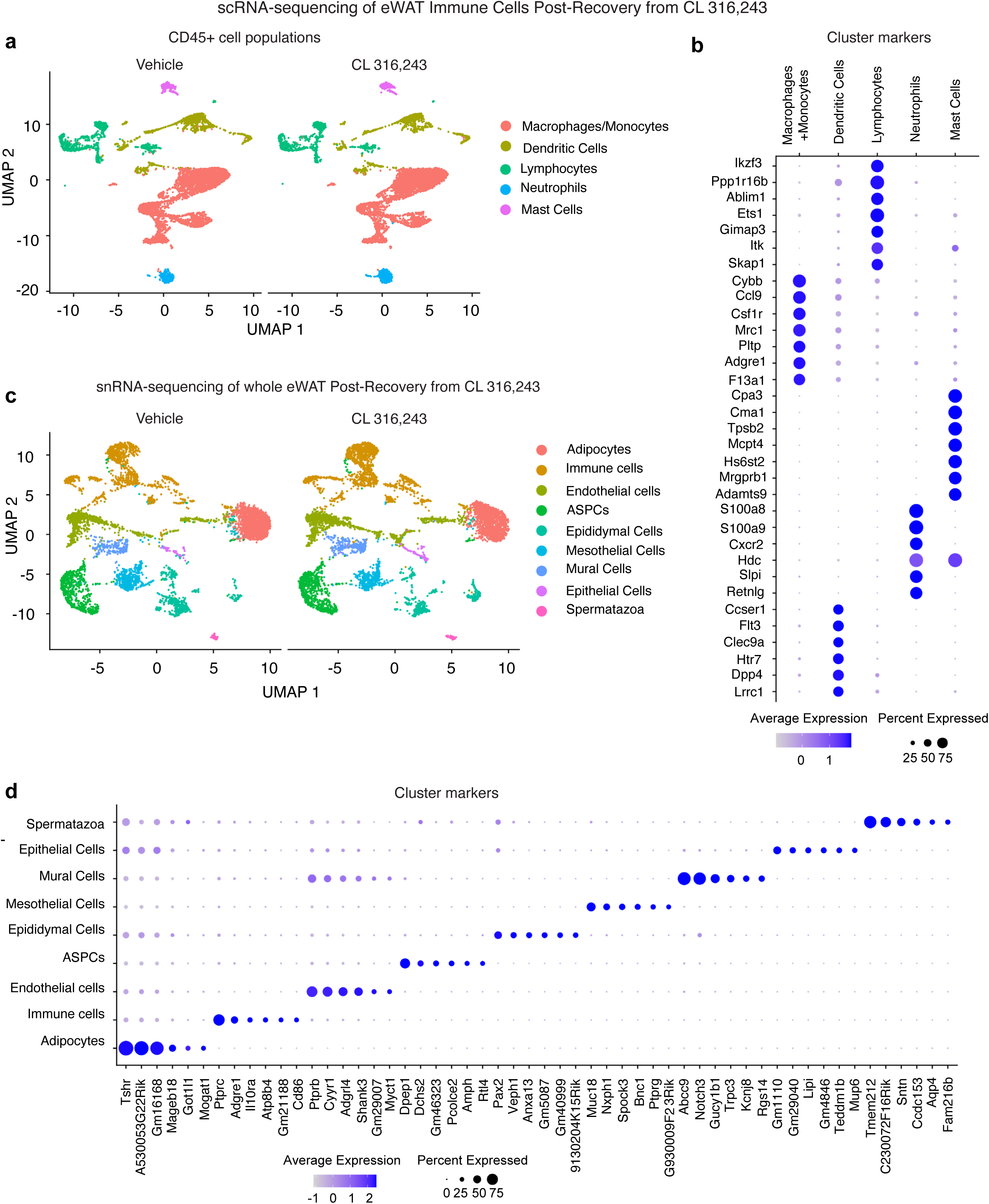
Cellular landscapes of eWAT after β3AR agonist withdrawal. **a)** Integrated Uniform Manifold Approximation and Projection (UMAP) of 14,851 CD45+ cell transcriptomes obtained from pooled eWAT depots of 4 mice previously treated with vehicle or 4 mice previously treated with CL 316,243. Data were filtered with the criteria of having more than 500 unique molecular identifiers (UMIs) per gene and more than 250 genes that consisting less than 10% in cells. The median number of UMIs detected per cell was 3,484 in the Vehicle group and 3,544 in the CL group. The mean reads per cell was 91,408 in the Vehicle group and 93,425 in the CL. **b)** Bubble plot of gene expression defining indicated immune cell types. **c)** UMAP of 12,504 nuclei transcriptomes obtained from pooled eWAT depots of 4 mice previously treated with vehicle or 4 mice previously treated with CL 316,243. Data were filtered with the criteria of having more than 500 unique molecular identifiers (UMIs) per gene and more than 250 genes that consisting less than 10% in cells. The median number of UMIs detected per cell was 1,882 in the Vehicle group and 1,810 in the CL group. The mean reads per cell was 28,820 in the Vehicle group and 18,354 in the CL group. **d)** Bubble plot of gene expression defining indicated eWAT cell types. Full datasets for panels b,d are in Supplementary Tables 13,29

**Extended Data Figure 3:**
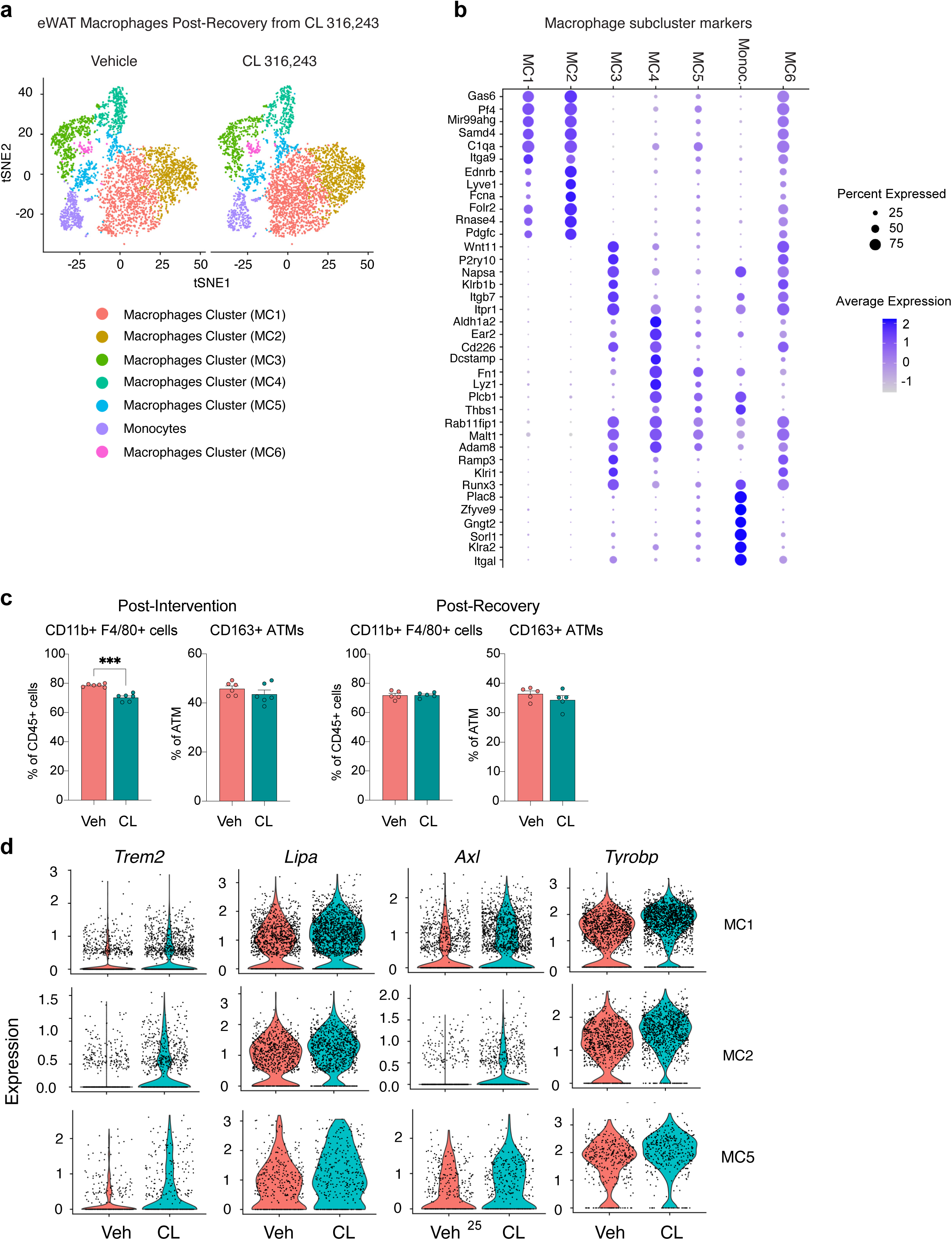
Macrophage transcriptional remodeling in eWAT following β3AR stimulation and withdrawal. **a)** UMAP of 8,313 eWAT macrophages/monocytes in vehicle-treated or CL-trained mice (Post-Recovery) (sub-clustered from CD45+ single cell RNA-seq shown in Extended Data Fig. 2). **b)** Bubble plot of gene expression defining indicated macrophage/monocyte subclusters. Full dataset of cluster markers is in Supplementary Tables 15. **c)** Frequency of total macrophages (CD45+ CD11b+ F4/80+) and CD163+ perivascular macrophages in eWAT following two weeks of vehicle or CL 316,243 treatment (Post-Intervention) and four weeks after cessation of the intervention (Post-Recovery). Bars represent mean + SEM. Each data point represents a unique animal. Exact measurements and statistical data are provided in Source Data Extended Data Figure 3. **d)** Violin plot of indicated genes associated with efferocytosis and lipid associated macrophages. Full datasets of differentially expressed genes (CL vs Veh.) and pathway analyses are in Supplementary Tables 14-28.

**Supplementary Figure 1:**
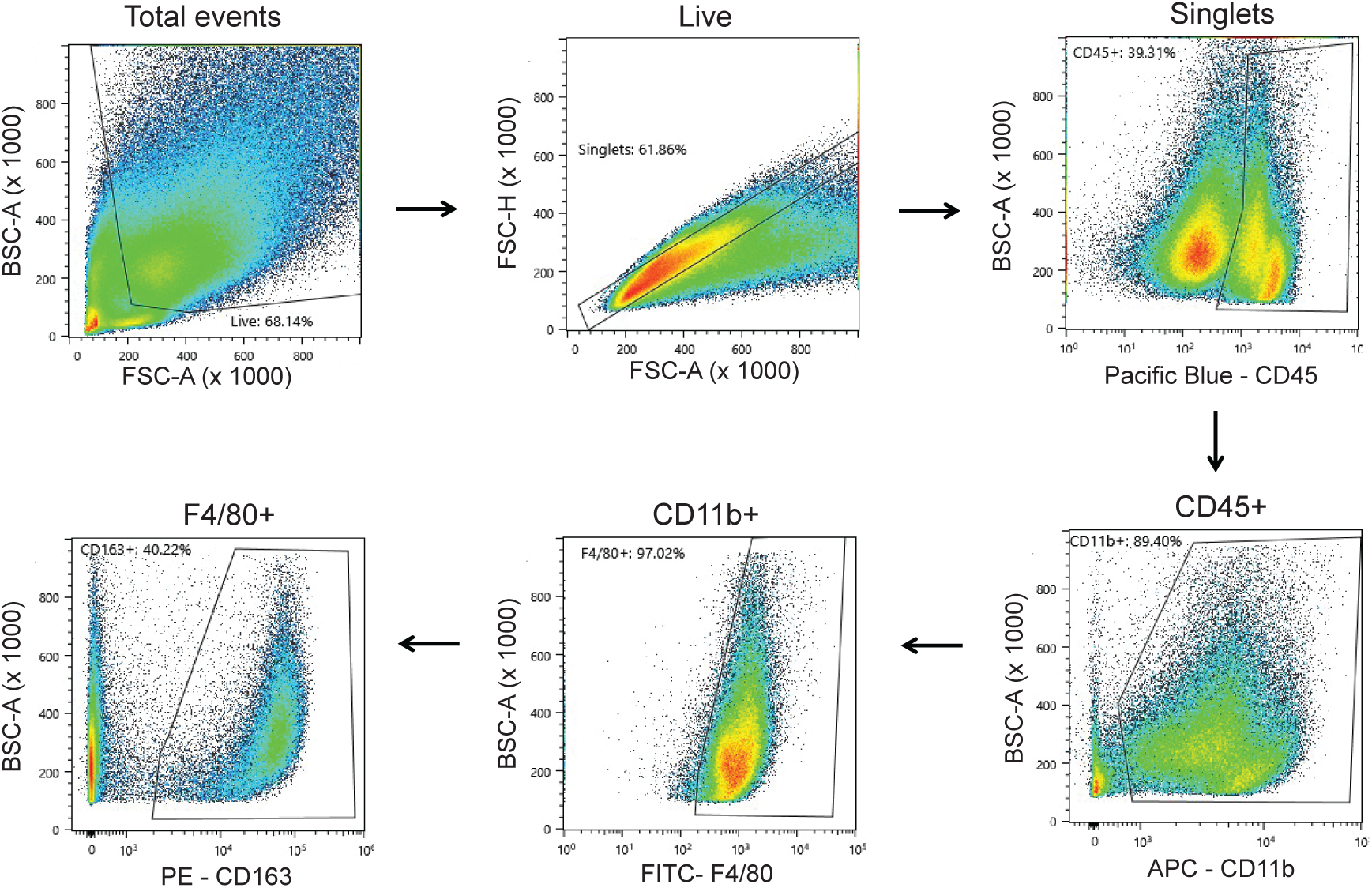
Flow cytometry gating strategy for the identification and quantification of adipose tissue macrophages (ATMs). (Related to Extended Data Figure 3). Live cells/single cells were gated from the stromal vascular fraction of collagenase-digested eWAT. Hematopoietic lineage cells (CD45+) were selected from live/single cells of the stromal vascular fraction of collagenase digested eWAT. Then total ATMs were identified from the CD45+ cells based on CD11b and F4/80 expression (CD45+ CD11b+ F4/80+). Perivascular macrophages were identified amongst ATMs based CD163 expression (CD163+).

**Extended Data Figure 4.**
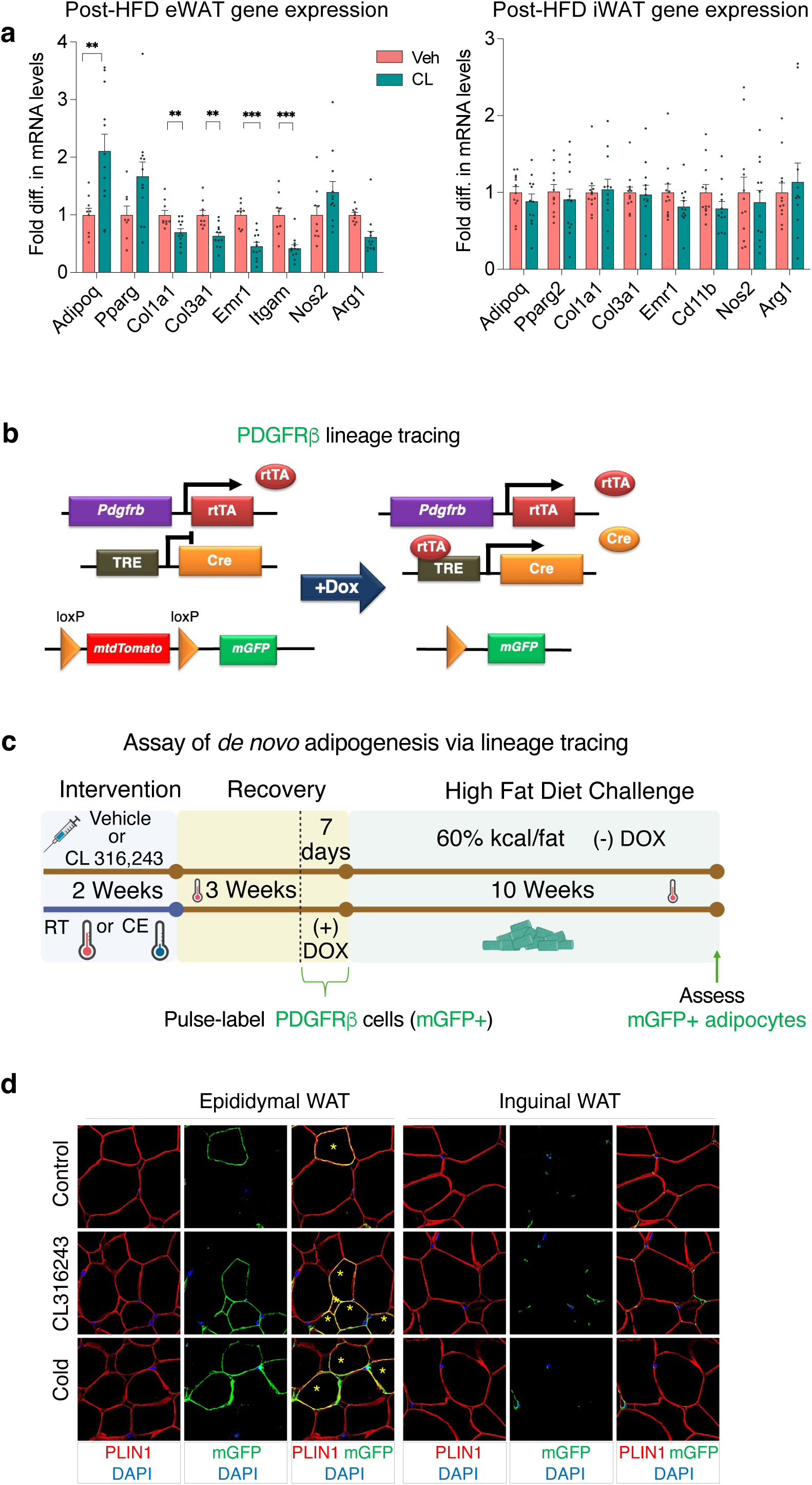
Prior cold exposure or β3AR agonism enhances future capacity for de novo adipogenesis during high-fat diet challenge. **a)** mRNA levels of indicated transcripts in whole eWAT or iWAT of control and CL-trained animals after 10 weeks of HFD feeding. n=11 mice per group. * denotes p<0.05, ** denotes p<0.01 by unpaired t-test. **b)** Pulse-Chase lineage tracing strategy: PDGFRβ+ adipose progenitor cells were labeled with mGFP expression in male *Pdgfrb*^rtTA^; *TRE*-CRE; *Rosa26R*^mT/mG^ mice with the addition of doxycycline (DOX) during the last week of recovery following CL 316,243 treatment. Animals were then placed on HFD (no DOX) (“Chase”) for 10 weeks prior to harvest. **c)** Representative confocal immunofluorescence images of PERILIPIN and mGFP expression in eWAT and iWAT after HFD feeding. mGFP+ PERILIPIN+ adipocytes represent newly formed adipocytes.

**Extended Data Figure 5.**
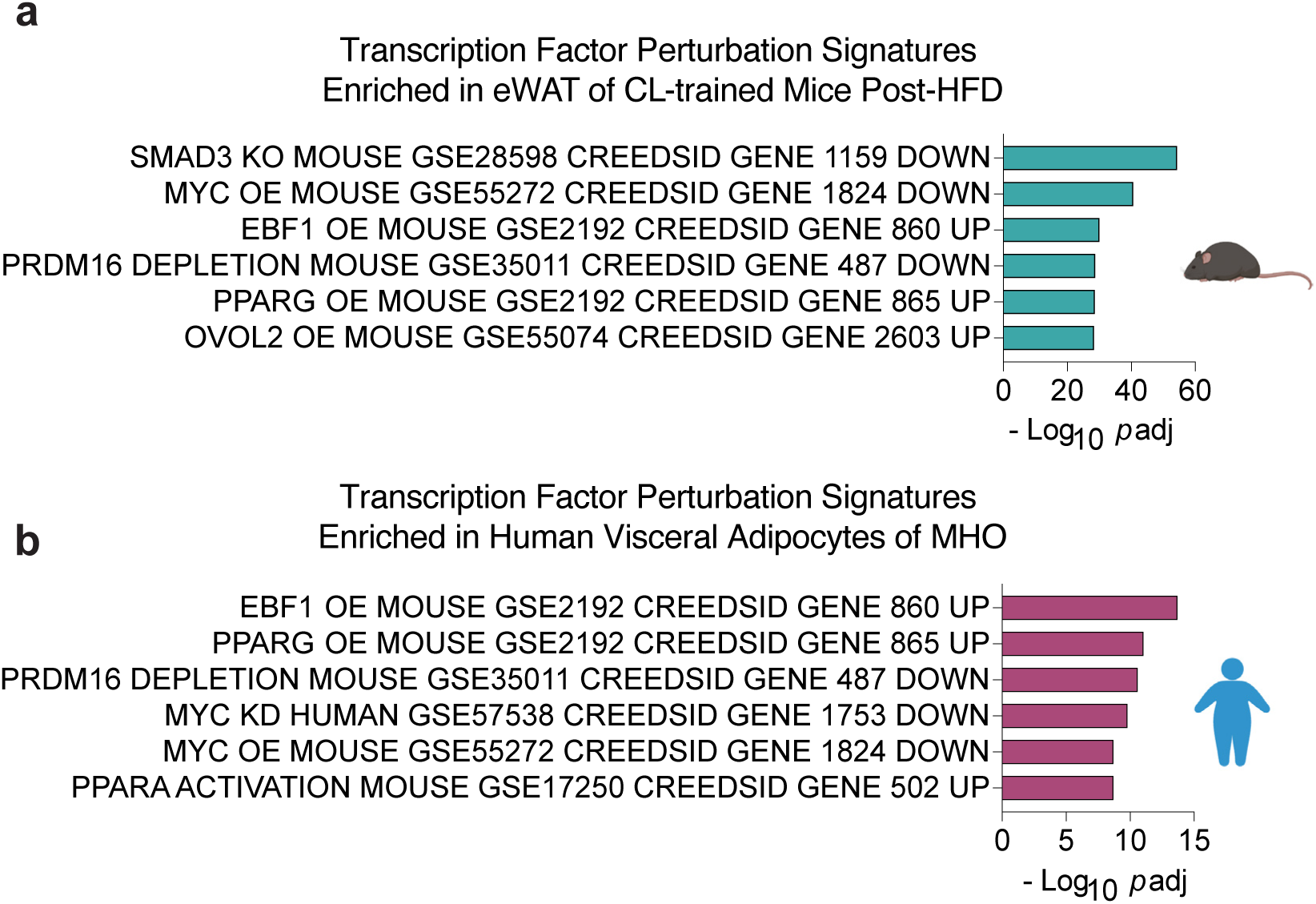
Adaptive adipose remodeling is linked to gene signatures of EBF, PPARγ, and PRDM16 activity. **a)** Top experimentally defined gene signatures of indicted transcription factor perturbations overrepresented (p adj. <0.05) in eWAT of CL-trained mice following high fat diet feeding (Post-HFD). **b)** Top experimentally defined gene signatures of indicted transcription factor perturbations enriched (p adj. <0.05) in visceral adipocytes of individuals with MHO compared to BMI-matched individuals with MUO. EnrichR analysis was performed using list of differentially expressed genes (MHO vs. MUO) described in Reinisch et al. Complete EnrichR analysis for panels a,b can be found in Supplementary Tables 38,40.

## References

1 Sakers, A., De Siqueira, M. K., Seale, P. & Villanueva, C. J. Adipose-tissue plasticity in health and disease. Cell 185, 419–446 (2022). 10.1016/j.cell.2021.12.016

2 Petersen, M. C. et al. Cardiometabolic characteristics of people with metabolically healthy and unhealthy obesity. Cell Metab 36, 745–761 e745 (2024). 10.1016/j.cmet.2024.03.002

3 Smith, G. I., Mittendorfer, B. & Klein, S. Metabolically healthy obesity: facts and fantasies. J Clin Invest 129, 3978–3989 (2019). 10.1172/JCI129186

4 Kim, S. M. et al. Loss of white adipose hyperplastic potential is associated with enhanced susceptibility to insulin resistance. Cell Metab 20, 1049–1058 (2014). 10.1016/j.cmet.2014.10.010

5 Kloting, N. et al. Insulin-sensitive obesity. Am J Physiol Endocrinol Metab 299, E506–515 (2010). 10.1152/ajpendo.00586.2009

6 Vishvanath, L. & Gupta, R. K. Contribution of adipogenesis to healthy adipose tissue expansion in obesity. J Clin Invest 129, 4022–4031 (2019). 10.1172/JCI129191

7 Shao, M. et al. De novo adipocyte differentiation from Pdgfrbeta(+) preadipocytes protects against pathologic visceral adipose expansion in obesity. Nat Commun 9, 890 (2018). 10.1038/s41467-018-03196-x

8 Crewe, C., An, Y. A. & Scherer, P. E. The ominous triad of adipose tissue dysfunction: inflammation, fibrosis, and impaired angiogenesis. J Clin Invest 127, 74–82 (2017). 10.1172/JCI88883

9 Divoux, A. et al. Fibrosis in human adipose tissue: composition, distribution, and link with lipid metabolism and fat mass loss. Diabetes 59, 2817–2825 (2010). 10.2337/db10-0585

10 Zhang, Q. et al. Distinct functional properties of murine perinatal and adult adipose progenitor subpopulations. Nat Metab 4, 1055–1070 (2022). 10.1038/s42255-022-00613-w

11 Cohen, P. & Kajimura, S. The cellular and functional complexity of thermogenic fat. Nat Rev Mol Cell Biol 22, 393–409 (2021). 10.1038/s41580-021-00350-0

12 Chouchani, E. T., Kazak, L. & Spiegelman, B. M. New Advances in Adaptive Thermogenesis: UCP1 and Beyond. Cell Metab 29, 27–37 (2019). 10.1016/j.cmet.2018.11.002

13 Yoshida, T., Sakane, N., Wakabayashi, Y., Umekawa, T. & Kondo, M. Anti-obesity and anti-diabetic effects of CL 316,243, a highly specific beta 3-adrenoceptor agonist, in yellow KK mice. Life Sci 54, 491–498 (1994). 10.1016/0024-3205(94)00408-0

14 Liu, X., Perusse, F. & Bukowiecki, L. J. Mechanisms of the antidiabetic effects of the beta 3-adrenergic agonist CL-316243 in obese Zucker-ZDF rats. Am J Physiol 274, R1212–1219 (1998). 10.1152/ajpregu.1998.274.5.R1212

15 Rosenwald, M., Perdikari, A., Rulicke, T. & Wolfrum, C. Bi-directional interconversion of brite and white adipocytes. Nat Cell Biol 15, 659–667 (2013). 10.1038/ncb2740

16 Roh, H. C. et al. Warming Induces Significant Reprogramming of Beige, but Not Brown, Adipocyte Cellular Identity. Cell Metab 27, 1121–1137 e1125 (2018). 10.1016/j.cmet.2018.03.005

17 Shao, M. et al. Cellular Origins of Beige Fat Cells Revisited. Diabetes 68, 1874–1885 (2019). 10.2337/db19-0308

18 Inoue, S. I. et al. Short-term cold exposure induces persistent epigenomic memory in brown fat. Cell Metab 36, 1764–1778 e1769 (2024). 10.1016/j.cmet.2024.05.011

19 Benvie, A. M. et al. Age-dependent Pdgfrbeta signaling drives adipocyte progenitor dysfunction to alter the beige adipogenic niche in male mice. Nat Commun 14, 1806 (2023). 10.1038/s41467-023-37386-z

20 Jaitin, D. A. et al. Lipid-Associated Macrophages Control Metabolic Homeostasis in a Trem2-Dependent Manner. Cell 178, 686–698 e614 (2019). 10.1016/j.cell.2019.05.054

21 Chavakis, T., Alexaki, V. I. & Ferrante, A. W., Jr. Macrophage function in adipose tissue homeostasis and metabolic inflammation. Nat Immunol 24, 757–766 (2023). 10.1038/s41590-023-01479-0

22 Sárvári, A. K. et al. Plasticity of Epididymal Adipose Tissue in Response to Diet-Induced Obesity at Single-Nucleus Resolution. Cell Metab 33, 437–453.e435 (2021). 10.1016/j.cmet.2020.12.004

23 Kuleshov, M. V. et al. Enrichr: a comprehensive gene set enrichment analysis web server 2016 update. Nucleic Acids Res 44, W90–97 (2016). 10.1093/nar/gkw377

24 Khan, T. et al. Metabolic dysregulation and adipose tissue fibrosis: role of collagen VI. Mol Cell Biol 29, 1575–1591 (2009). 10.1128/MCB.01300-08

25 Johannsen, D. L. et al. Effect of 8 weeks of overfeeding on ectopic fat deposition and insulin sensitivity: testing the “adipose tissue expandability” hypothesis. Diabetes Care 37, 2789–2797 (2014). 10.2337/dc14-0761

26 Shao, M. et al. Pathologic HIF1alpha signaling drives adipose progenitor dysfunction in obesity. Cell Stem Cell 28, 685–701 e687 (2021). 10.1016/j.stem.2020.12.008

27 Vishvanath, L. et al. Pdgfrbeta+ Mural Preadipocytes Contribute to Adipocyte Hyperplasia Induced by High-Fat-Diet Feeding and Prolonged Cold Exposure in Adult Mice. Cell Metab 23, 350–359 (2016). 10.1016/j.cmet.2015.10.018

28 Reinisch, I. et al. Unveiling adipose populations linked to metabolic health in obesity. Cell Metab 37, 640–655 e644 (2025). 10.1016/j.cmet.2024.11.006

29 Efthymiou, V. et al. Single-Nucleus Analysis of Human White Adipose Tissue Reveals Adipocyte Subsets with Distinct Metabolic Profiles. bioRxiv (2025). 10.1101/2025.09.14.673351

30 Wander, G. S. & Bansal, M. Legacy effect in medicine-the expanding horizon! Indian Heart J 70, 769–771 (2018). 10.1016/j.ihj.2018.12.001

31 Lundgren, P. et al. A subpopulation of lipogenic brown adipocytes drives thermogenic memory. Nat Metab 5, 1691–1705 (2023). 10.1038/s42255-023-00893-w

32 Sun, W. et al. Cold-induced epigenetic programming of the sperm enhances brown adipose tissue activity in the offspring. Nat Med 24, 1372–1383 (2018). 10.1038/s41591-018-0102-y

33 van Beek, S., Hashim, D., Bengtsson, T. & Hoeks, J. Physiological and molecular mechanisms of cold-induced improvements in glucose homeostasis in humans beyond brown adipose tissue. Int J Obes (Lond) 47, 338–347 (2023). 10.1038/s41366-023-01270-z

34 O’Mara, A. E. et al. Chronic mirabegron treatment increases human brown fat, HDL cholesterol, and insulin sensitivity. J Clin Invest 130, 2209–2219 (2020). 10.1172/JCI131126

35 Speakman, J. R. & Heidari-Bakavoli, S. Type 2 diabetes, but not obesity, prevalence is positively associated with ambient temperature. Sci Rep 6, 30409 (2016). 10.1038/srep30409

36 Johnson, J. L., Slentz, C. A., Ross, L. M., Huffman, K. M. & Kraus, W. E. Ten-Year Legacy Effects of Three Eight-Month Exercise Training Programs on Cardiometabolic Health Parameters. Front Physiol 10, 452 (2019). 10.3389/fphys.2019.00452

37 Jeffery, E. et al. The Adipose Tissue Microenvironment Regulates Depot-Specific Adipogenesis in Obesity. Cell Metab 24, 142–150 (2016). 10.1016/j.cmet.2016.05.012

38 Shan, B. et al. Multilayered omics reveal sex- and depot-dependent adipose progenitor cell heterogeneity. Cell Metab 34, 783–799 e787 (2022). 10.1016/j.cmet.2022.03.012

39 Hepler, C. et al. Directing visceral white adipocyte precursors to a thermogenic adipocyte fate improves insulin sensitivity in obese mice. Elife 6 (2017). 10.7554/eLife.27669

40 Seale, P. et al. Prdm16 determines the thermogenic program of subcutaneous white adipose tissue in mice. J Clin Invest 121, 96–105 (2011). 10.1172/JCI44271

41 Griffin, M. J. et al. Early B-cell factor-1 (EBF1) is a key regulator of metabolic and inflammatory signaling pathways in mature adipocytes. J Biol Chem 288, 35925–35939 (2013). 10.1074/jbc.M113.491936

42 Gao, H. et al. Early B cell factor 1 regulates adipocyte morphology and lipolysis in white adipose tissue. Cell Metab 19, 981–992 (2014). 10.1016/j.cmet.2014.03.032

43 Petrus, P. et al. Low early B-cell factor 1 (EBF1) activity in human subcutaneous adipose tissue is linked to a pernicious metabolic profile. Diabetes Metab 41, 509–512 (2015). 10.1016/j.diabet.2015.02.004

44 Rahbani, J. F. et al. ADRA1A-Galpha(q) signalling potentiates adipocyte thermogenesis through CKB and TNAP. Nat Metab 4, 1459–1473 (2022). 10.1038/s42255-022-00667-w

45 Dobin, A. et al. STAR: ultrafast universal RNA-seq aligner. Bioinformatics 29, 15–21 (2013). 10.1093/bioinformatics/bts635

46 Liao, Y., Smyth, G. K. & Shi, W. featureCounts: an efficient general purpose program for assigning sequence reads to genomic features. Bioinformatics 30, 923–930 (2014). 10.1093/bioinformatics/btt656

47 Love, M. I., Huber, W. & Anders, S. Moderated estimation of fold change and dispersion for RNA-seq data with DESeq2. Genome Biol 15, 550 (2014). 10.1186/s13059-014-0550-8

48 So, J. et al. Robust single-nucleus RNA sequencing reveals depot-specific cell population dynamics in adipose tissue remodeling during obesity. Elife 13 (2025). 10.7554/eLife.97981

49 Battenberg, K. et al. A flexible cross-platform single-cell data processing pipeline. Nat Commun 13, 6847 (2022). 10.1038/s41467-022-34681-z

50 Young, M. D. & Behjati, S. SoupX removes ambient RNA contamination from droplet-based single-cell RNA sequencing data. Gigascience 9 (2020). 10.1093/gigascience/giaa151

51 Hao, Y. et al. Dictionary learning for integrative, multimodal and scalable single-cell analysis. Nat Biotechnol 42, 293–304 (2024). 10.1038/s41587-023-01767-y

52 Tarazona, S. et al. Data quality aware analysis of differential expression in RNA-seq with NOISeq R/Bioc package. Nucleic Acids Res 43, e140 (2015). 10.1093/nar/gkv711

53 Jin, S. et al. Inference and analysis of cell-cell communication using CellChat. Nat Commun 12, 1088 (2021). 10.1038/s41467-021-21246-9

